# MiR-126 impairs the intestinal barrier function via inhibiting S1PR2 mediated activation of PI3K/AKT signaling pathway

**DOI:** 10.1101/110338

**Authors:** Tanzhou Chen, Haibo Xue, Ruoyang Lin, Zhiming Huang

## Abstract

**Background:** Aberrant expression of miRNAs was a critical element in the pathogenesis of inflammatory bowel disease (IBD). This study aimed to explore the involvement and mechanism of miR-126 in IBD.

**Methods:** In this study, the endogenous expressions of miR-126, S1PR2 and S1P in the pathological tissues of patients with IBD were detected using qRT-PCR and western blot assay, respectively. The luciferase reporter gene assay was performed to confirm the targeting regulatory relation between miR-126 and S1PR2. The transendothelial electrical resistance assay was used to measured the value of TEER.

**Results:** The expressions of miR-126, S1PR2 and S1P in the pathological tissues of IBD patients were significantly higher than that of the control group. Moreover, miR-126 overexpression contributed to intestinal mucosal barrier dysfunction *in vitro.* S1PR2 was a direct target of miR-126, and S1PR2 expression was negatively regulated by miR-126 in Caco-2 cells. However, S1PR2 activated by S1P had the protection effect for the integrity and permeability of intestinal mucosal barrier via a PI3K/Akt dependent mechanism. MiR-126 silencing possessed obvious protective effects on the intestinal barrier function, but these effects could be reversed by JTE-013 or LY294002.

**Conclusion:** MiR-126 down-regulated S1PR2 and then prevented the activation of PI3K/AKT signaling pathway, which ultimately could damage intestinal mucosal barrier function.

## Introduction

Inflammatory bowel disease (IBD) is a kind of chronic intestinal inflammatory disease, including ulcerative colitis (UC) and Crohn’s disease (CD)[1], but the etiology of IBD is still uncertain as yet. The prevalence rate of IBD was high in many western countries, with a morbidity of 1-2/1000. China had showing a obvious increased incidence of IBD for nearly a decade, which was the 10-20 times of that of ten years ago. In view of the serious harm of IBD to human health and the high deterioration rate, more and more researchers have interests in its pathogenesis and treatment. The mechanism was hypothesized as that IBD was caused by the interaction of multiple factors, such as environment, heredity, infection and immunity [2].

MicroRNAs (miRNAs) are a class of non-coding RNA molecules, about 20-23 nucleotides long and identified as important transcriptional regulation factors[3]. MiRNAs can regulate the level of target molecules through several different mechanisms, including transcriptional and translational level and gene silencing through directly binding to the 3’UTR of target mRNA[4]. It has been confirmed that the abnormality of miRNAs expression was a critical element in the pathogenesis of human diseases. MiR-126 was found to significantly increase in active UC tissues and contribute to the development of UC via activation of NF-κB signaling pathway[5]. However, the concrete machanism of miR-126 in IBD needs to be discussed and explored further, which is the focus of this study.

Sphingosine-1-phosphate (S1P), a signaling sphingolipid of the cell membrane, could regulate a variety of physiological processes of cells, such as growth, apoptosis, proliferation and migration[6]. The extensive studies verified that S1P was involved in the pathophysiologic process of many diseases, as well as the pathogenesis of IBD[7]. S1P signaling was also served as an important mediator in the process of lymphocyte trafficking[8]. Several publications demonstrated that S1P was closely related to autoimmune disease through S1P receptors (S1PRs)[9]. In previous studies, S1P exerted its biological effects mainly through its receptors, S1P receptors 1–5 (S1PR1–5) belonged to S1PRs. The results of animal model experiments indicated that targeting S1P signaling might be a potential therapeutic method for IBD[10]. It is an important regulator of vascular endothelial cell permeability and fluid balance in vivo and is able to enhance endothelial barrier. Recent studies found that S1P could regulate the permeability of vascular endothelial cells and also enhance endothelial barrier function[11]. Moreover, S1PR2 had inhibition on the endothelial barrier function[12], but the role of S1PR2 in IBD has not been reported. Bioinformatics analysis showed the putative binding site of miR-126 and S1PR2, so we aimed to explore that whether miR-126 was involved in the intestinal mucosal barrier dysfunction via S1PR2.

## Materials and methods

### Patients

20 patients with colorectal neoplasms under the endoscopy were recruited, which were regarded as the model group. Normal intestinal mucosa tissues and another 5 patients with benign enteropathy were chosen as control. Our study was approved by the human ethics committee of The First Affiliated Hospital of Wenzhou Medical University.

### Cell line and culture

Caco-2 cells, which were the normal intestinal epithelial cell line, were purchased from iCellBiotechnology Co., Ltd (China). The cells were maintained in RPBM 1640 medium supplemented with 10% fetal calf serum under controlled temperature of 37 °C and a humidified 5% CO_2_.

### Luciferase reporter assay

According on Genebank online, the fragment of S1PR2 was obtained. The luciferase reporter plasmid was recombined with the 3’UTR of S1PR2. S1PR2-3’UTR vector and miR-126 mimic or miRNA were co-transfected into the cells used Lipofectamine 2000 transfection reagent (Invitrogen, USA) in accordance with the manufacture’s instruction. The measurement of the luciferase activity was used with Dual-Luciferase Reporter Assay System.

### Cell transfection

The miR-126 mimic, inhibitor and their negative control (NC) were synthesized by GENEWIZ Biotechnology Co., Ltd (Suzhou, China). Cells were planted in the 96-well plates and maintained for 24 h. Cell transfection was used by Lipofectamine 2000 reagent (Invitrogen) according to the manufacture’s instructions. The efficiency of transfection was examined by real-time PCR.

### Real-time PCR

Oligotex Direct mRNA Micro Kit (QIAGEN) was used to isolate total RNA from cells, then 1µL of RNA was quantitative. Equal amount of RNA was reverse transcribed for the synthesis of cDNA. Real-time PCR was performed with PCR Mix (SYBR Green I) on the Real-time PCR system (ABI 7900 HT). In this study, the primers were synthesized by GENEWIZ Biotechnology Co., Ltd (Suzhou, China). GAPDH was considered as internal control.

### Western blot

With the lysis in RIPA buffer (Solarbio) and centrifugation for 20 min, proteins were extracted from cells. Bradford method was used to detect the quality of the proteins. SDS-PAGE was used to separate the proteins with equal amount, the proteins were then transferred onto PVDF membrane, with the primary antibodies, which were purchased from Sigma (USA), against S1P, S1PR2, ZO-1and p-AKT at 4 °C for 24 h, then the membrane was incubated with the secondary antibodies at room temperature for another 1 h. β-actin acted as internal control.

### Intestinal epithelial barrier model construction and Trans-epithelial electrical resistance (TER) determination

Caco-2 cells were seeded in a Transwell system for monolayers, TER values were measured by an epithelial voltohmmeter ERS-2 (Merck Millipore). TER was measured until the similar values were occurred on three consecutive measurements. The values were calculated with Ω cm^2^. When the TER values of the cells reached at least 500 Ω cm^2^ at 20 d, showing the model was successfully established.

### Paracellular permeability measurement

After intestinal epithelial barrier model was successfully constructed, LPS was added in the Apical side of the model and cultured with the supplementary of 1 mg/mL FD-4 solution for 2 h. The paracellular permeability was measured by FD-4 flux. A Synergy H2 microplate reader (Bio Tek) was used for the determination of FD-4 signaling.

### Statistical analysis

Data were presented as means±SD by statistical analysis of ANOVA software combined with t-test. The value of P less than 0.05 was considered as statistically significant difference.

## Results

### MiR-126, S1PR2 and S1P with high expression in IBD

The expression of miR-126 in the pathological tissues of IBD patients (n=20) was significantly higher than that of the control group (about 2.8-fold) (n=5) (Figure 1 A). Moreover, the mRNA expression of S1PR2 was also up-regulated by 1.61-fold in the pathological tissues of IBD patients (n=20) (Figure 1 B), and it negatively correlated with miR-126 expression (Figure 1 C). And the protein levels of S1PR2 and S1P expression were significantly increased in the pathological tissues of IBD patients (n=20) (Figure 1 D&E).

**Figure 1.**
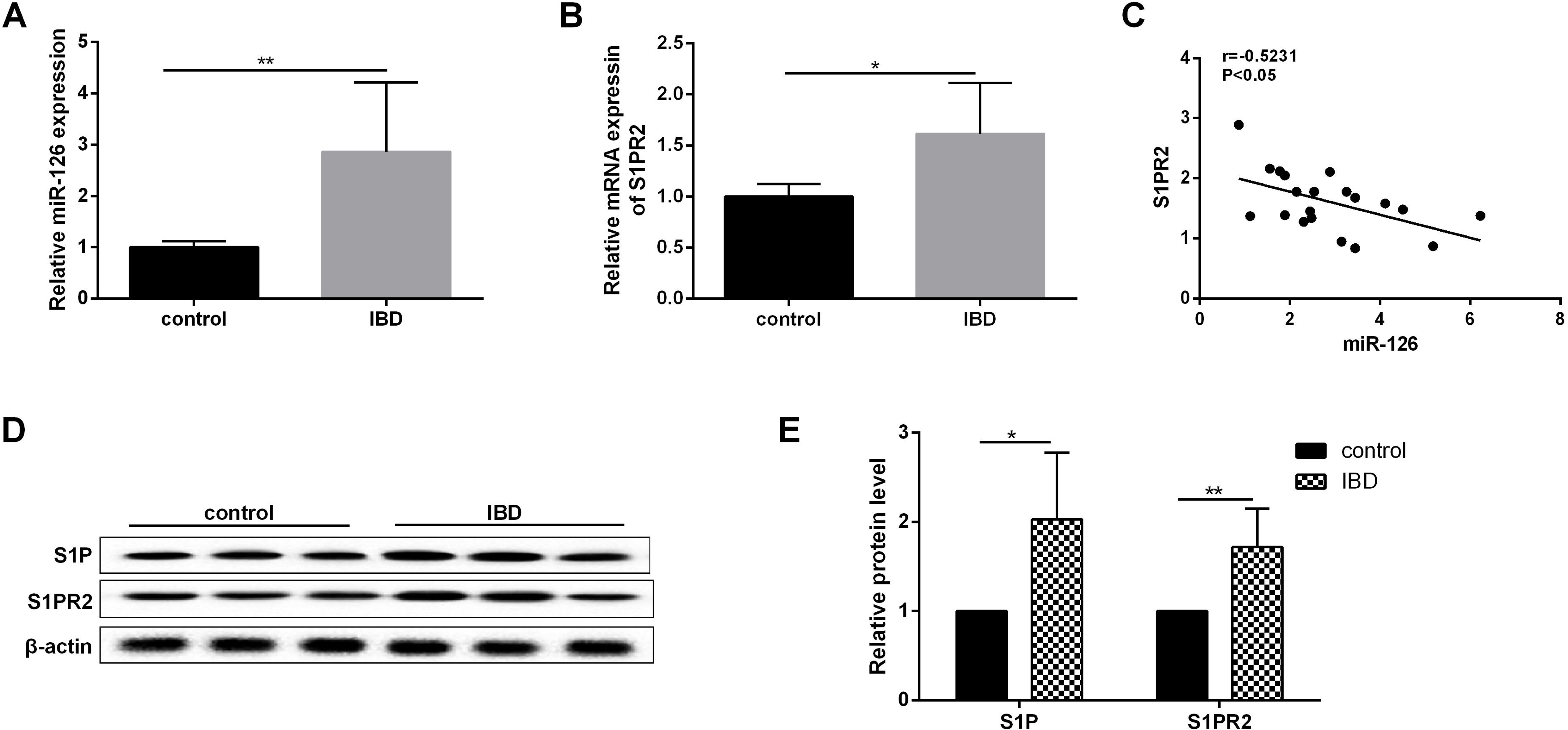
The endogenous expression of miR-126, S1PR2 and S1P in the pathological tissues of patients with IBD. The mRNA expressions of **(A)** miR-126 and **(B)** S1PR2 in the pathological tissues of IBD patients (n=20) were detected using qRT-PCR, the normal intestinal mucosal tissues (n=5) from patients at the remission phase of IBD as the control. **(C)** Analysed the correlation between the expression of miR-126 and S1PR2 in the pathological tissues of patients with IBD. **(D-E)** The protein expressions of S1PR2 and S1P in the pathological tissues and normal intestinal mucosal tissues were measured by western blot assay. *P<0.05,**P<0.01 vs. control.

### S1PR2 was a direct target of miR-126

The results of bioinfomatics analysis revealed that miR-126 directly bond at a site located within S1PR2 (Figure 2 A). The relative luciferase activity of S1PR2 was significantly suppressed by miR-126 mimic, but the relative luciferase activity was unaffected in miRNC treatment (Figure 2 A). The protein expression of S1PR2 was decreased with miR-126 mimic; conversely, S1PR2 protein was significantly elevated by miR-126 inhibitor (Figure 2 B).

**Figure 2.**
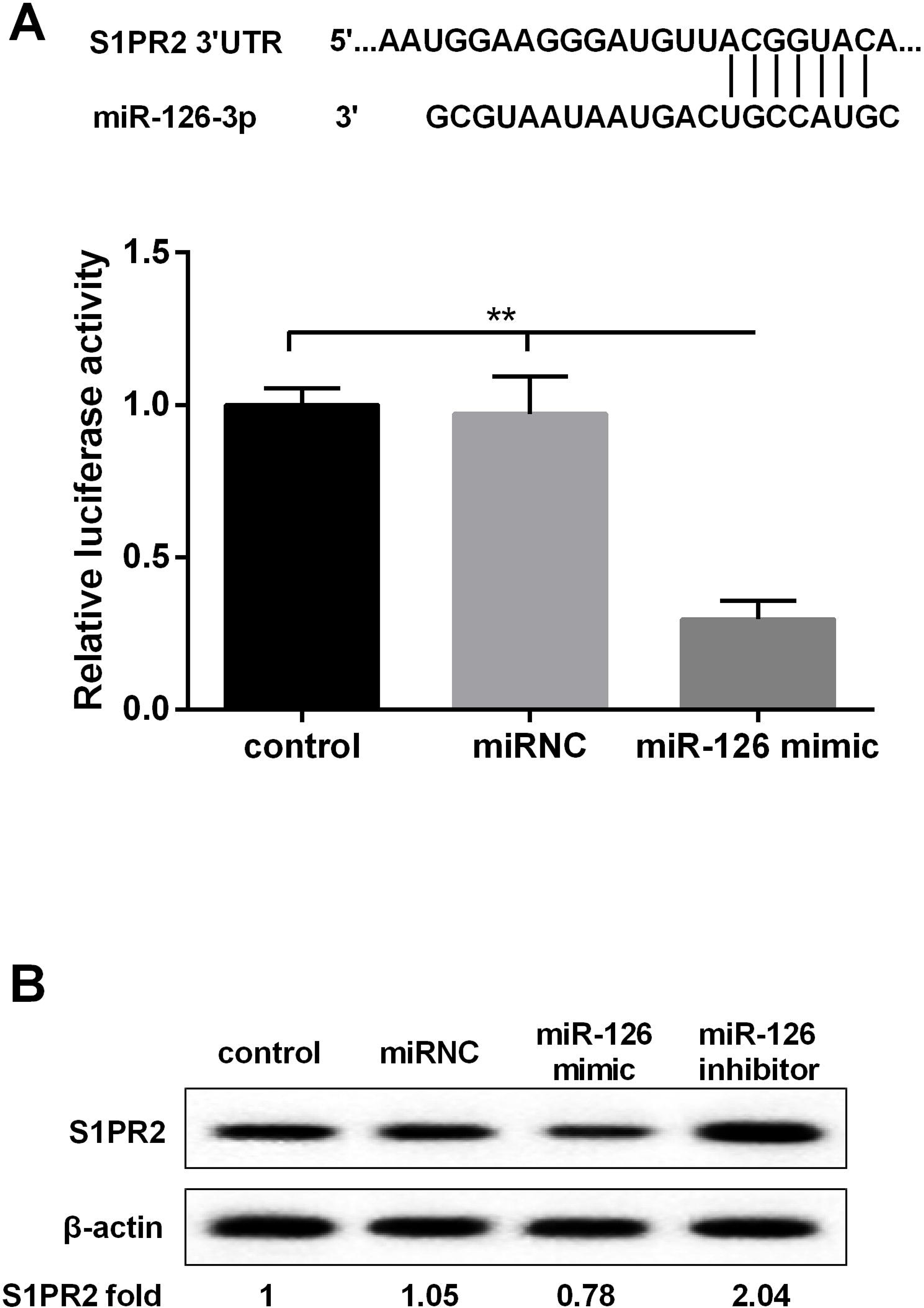
MiR-126 could inhibit S1PR2 expression by targeting to S1PR2. **(A)** The putative binding site of miR-126 and S1PR2 was exhibited and luciferase reporter gene assay was performed to confirm the targeting regulatory relation. **(B)** The effects of overexpression or knockdown of miR-126 on the expression of S1PR2 protein. **P<0.01 vs. control.

### MiR-126 overexpression contributed to intestinal mucosal barrier dysfunction

LPS was used for the induction of intestinal mucosal barrier dysfunction and caused continuous reduction of TEER in Caco-2 cells at 0h, 1h, 3h, 6h, 12h of LPS treatment (Figure 3 A). Moreover, S1P significantly inhibited the decline of TEER induced by LPS, but miR-126 mimic could largely release the effect of S1P on TEER (Figure 3 A). Furthermore, miR-126 mimic also significantly reversed the S1P-mediated decrease of intestinal paracellular permeability of Caco-2 cells (Figure 3 B). S1P enhanced the expression of S1PR2 and ZO-1 protein in Caco-2 cells treated by LPS, but this increase of S1PR2 and ZO-1 was completely reversed by miR-126 mimic (Figure 3 C).

**Figure 3.**
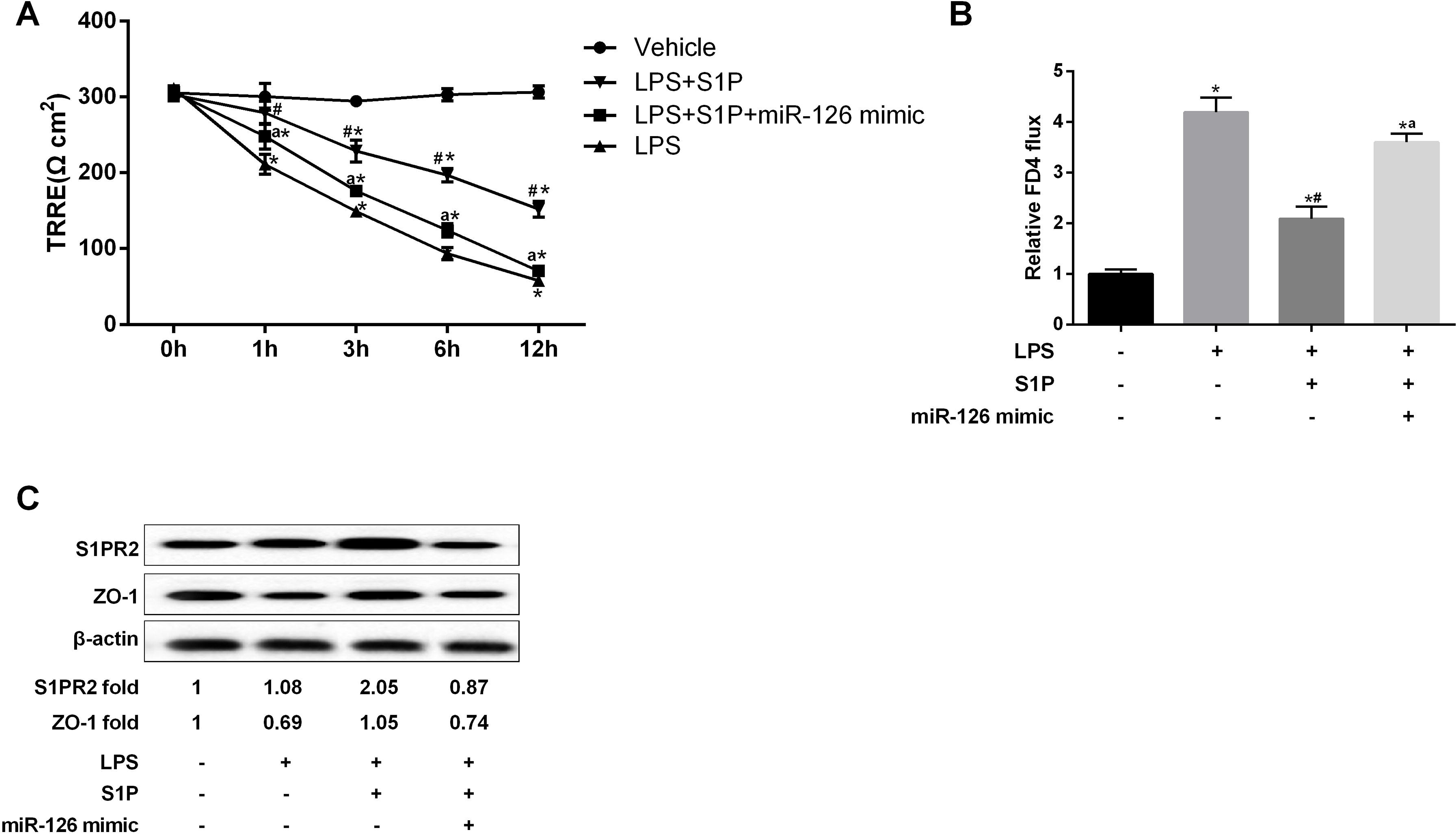
The effects of miR-126 overexpression on intestinal mucosal barrier dysfunction. Caco-2 cells were treated with ddH_2_O (vehicle group), LPS+S1P, LPS+S1P+miR-126 mimic, LPS, respectively. **(A)** The TEER and **(B)** intestinal paracellular permeability of Caco-2 cells were measured. **(C)** The protein expression of S1PR2 and ZO-1 was determined using western blot assay. *P<0.05 vs. Vehicle, ^#^P<0.05 vs. LPS, ^a^P<0.05 vs. LPS+S1P.

### S1P regulated intestinal barrier function via modulation of SP1R2

Si-SP1R2 or JTE-013, an antagonist of S1PR2, could largely release the effect of S1P on the TEER and intestinal paracellular permeability of Caco-2 cells; while CYM-5478, a prototypical agonist of SP1R2, magnified the effect of S1P as shown by increased TEER and decreased intestinal paracellular permeability *in vitro* (Figure 4 A&B). In addition, similar results were also observed in the protein expression of ZO-1, p-AKT and AKT. Si-SP1R2 or JTE-013 prominently attenuated the up-regulation of ZO-1 and p-AKT/AKT induced by S1P, but CYM-5478 further promoted the increase of ZO-1 and p-AKT/AKT (Figure 4 C&D&E).

**Figure 4.**
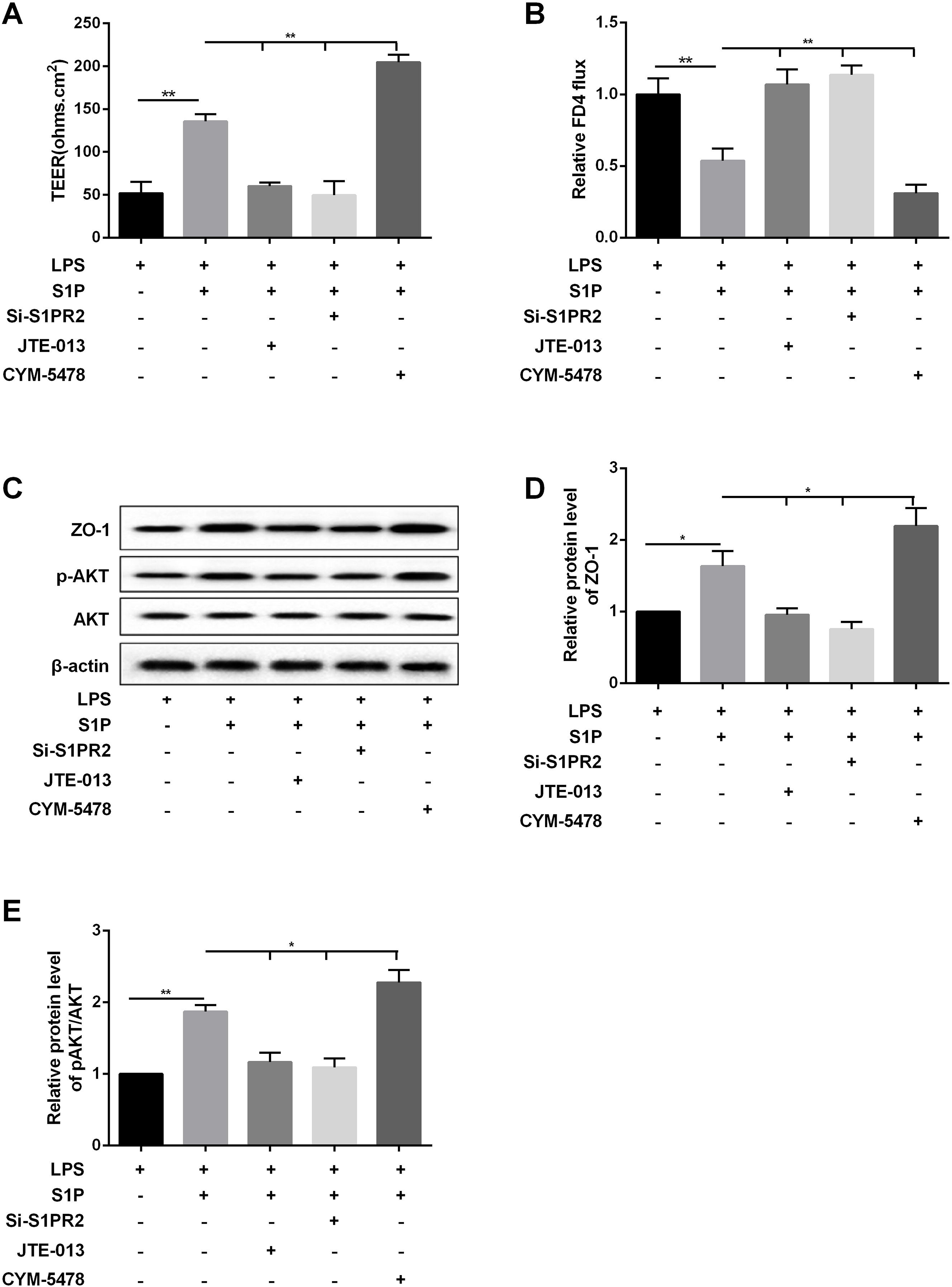
S1P regulated intestinal barrier function via SP1R2 expression. Caco-2 cells were treated with LPS, LPS+S1P, LP S+S1P+JTE-013, LP S+S1 P+si-SP1R2, LPS+S1P+CYM-5478, respectively. JTE-013 is the antagonist of S1PR2, and CYM-5478 is a prototypical agonist of SP1R2. **(A)** The TEER and **(B)** intestinal paracellular permeability of Caco-2 cells were measured. **(C-E)** The protein expression of ZO-1, p-AKT and AKT was determined using western blot assay. *P<0.05, **P<0.01.

### S1P/S1PR2 regulated tight junction protein ZO-1 through PI3K/AKT signaling pathway

LY294002 treatment decreased p-AKT/AKT as well as the protein expression of ZO-1 (Figure 5 A). CYM-5478 had the obvious effect of promoting p-AKT and ZO-1 at the protein level, but LY294002 almost completely inhibited the increase of p-AKT and ZO-1 protein induced by CYM-5478 (Figure 5 B).

**Figure 5.**
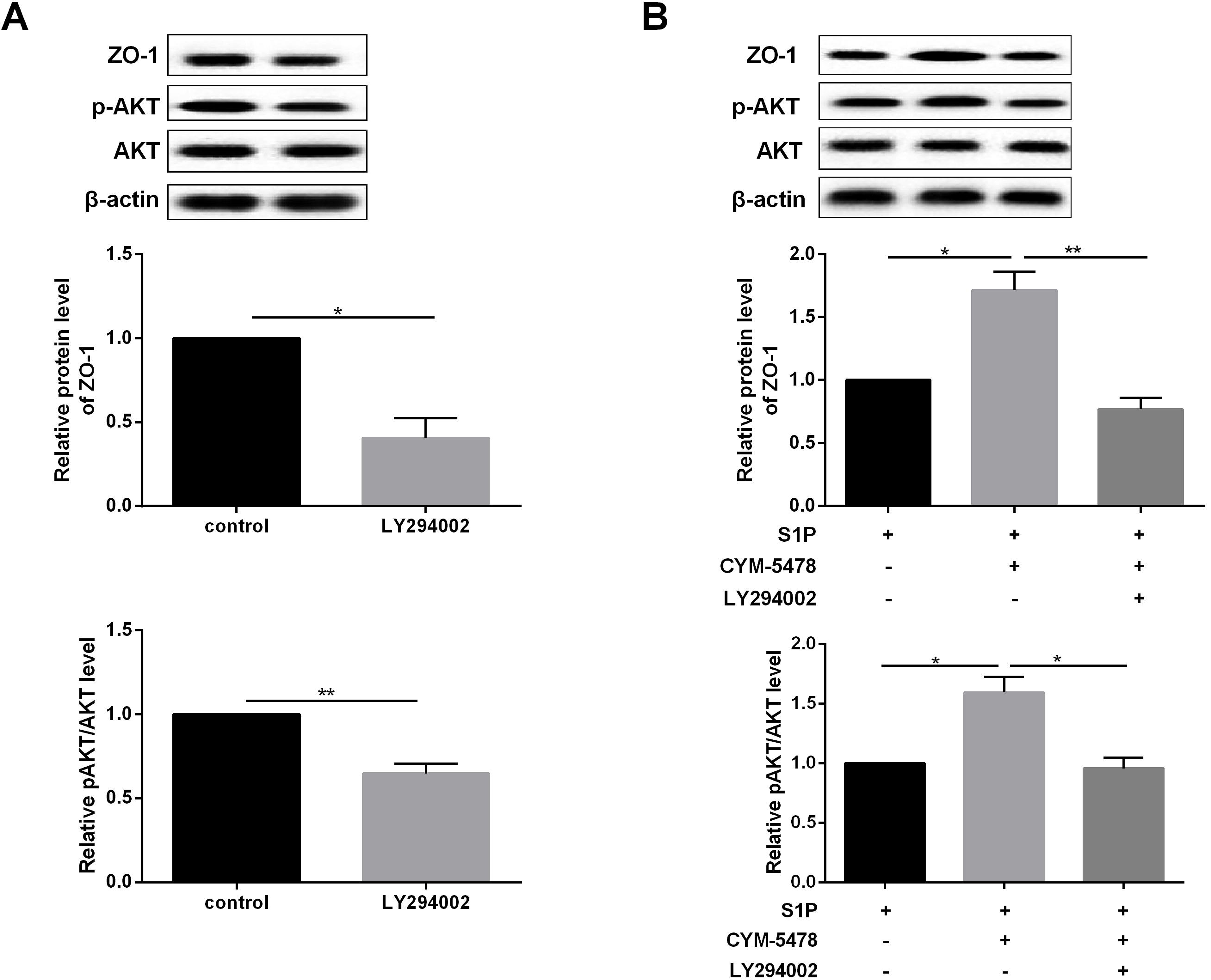
S1P/S1PR2 regulated tight junction protein ZO-1 through PI3K/AKT signaling pathway. **(A)** After treatment by LY294002, the protein expression of ZO-1, p-AKT and AKT in Caco-2 cells was measured by western blot assay. **(B)** After treatment with CYM-5478 or CYM-5478+LY294002, the protein expression of ZO-1, p-AKT and AKT in Caco-2 cells was measured by western blot assay. *P<0.05, **P<0.01.

### The role of miR-126 in the regulation mechanism of S1P/S1PR2/PI3K/AKT signaling pathway for intestinal barrier function

TEER of Caco-2 cells was elevated by miR-126 inhibitor, but JTE-013 or LY294002 could down-regulate this increase (Figure 6 A). JTE-013 or LY294002 also reversed the miR-126 inhibitor-mediated repression of intestinal paracellular permeability of Caco-2 cells (Figure 6 B), and eliminated the effect of miR-126 inhibitor on the protein expression of S1PR2, p-AKT, AKT and ZO-1 (Figure 6 C).

**Figure 6.**
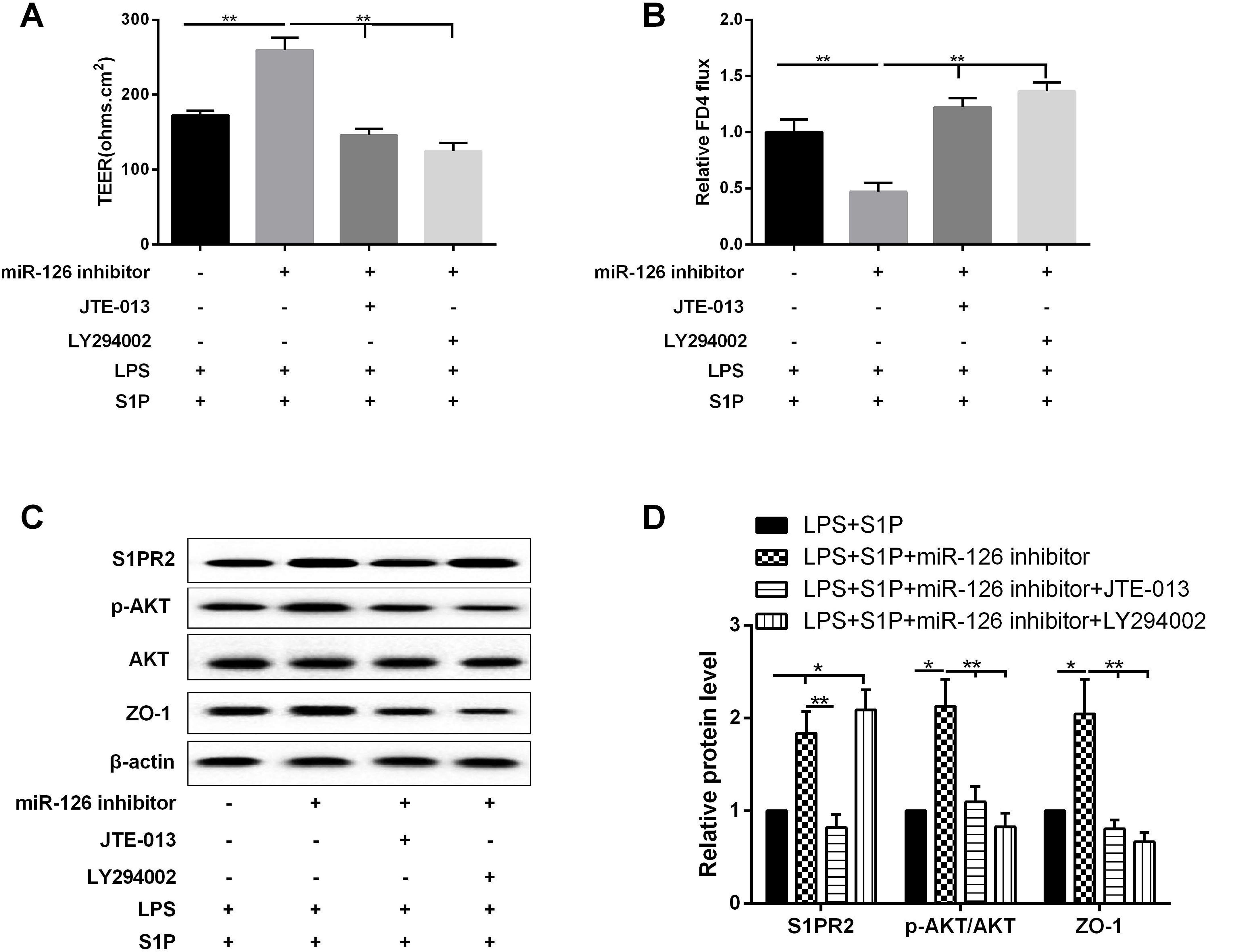
The role of miR-126 in the regulation mechanism of S1P/S1PR2/PI3K/AKT signaling pathway for intestinal barrier function. Caco-2 cells were treated with LPS+S1P, LPS+S1P+miR-126 inhibitor, LPS+S1P+miR-126 inhibitor+JTE-013, LPS+S1P+miR-126 inhibitor+LY294002, respectively. **(A)** The TEER and **(B)** intestinal paracellular permeability of Caco-2 cells were measured in different treatment groups. **(C)** The protein expression of S1PR2, p-AKT, AKT and ZO-1 in the cells was detected using western blot assay. *P<0.05, **P<0.01.

## Discussion

MiR-126 was convinced to be participated in the genesis and development of ulcerative colitis (UC), but its functional role and mechanism in inflammatory bowel disease (IBD) have not been known well. In the current study, we found that the high expression of miR-126 in IBD had potential destructive effects on the intestinal barrier function via targeted modulation of S1PR2. Furthermore, we also demonstrated that the regulatory mechanism of miR-126/S1PR2 axis was associated with PI3K/AKT signaling pathway.

Our results identified that miR-126 was up-regulated in the pathological tissues of IBD patients, which was consistent with the previous findings. It was also observed that miR-126 significantly increased in active UC group and contributed to the pathogenesis of UC[5]. Altogether, these results indicated that miR-126 may serve as a promotion factor for the progress of IBD. Furthermore, we also found that S1P and its receptor, S1PR2 were increased in IBD patients. S1P, a membrane sphingolipid, can enhance the intestinal epithelial barrier function via activating S1PRs. Additionally, S1PR2 was activated by S1P and promoted the proliferation of cholangiocarcinoma cells[13]. It has been reported that S1PR2 can regulate the migration, proliferation and differentiation of mesenchymal stem cells[14]. Our results suggested that S1PR2 also played an important role in IBD. Further analysis showed that there was a negative correlation between miR-126 and S1PR2 expression in the pathological tissues of IBD, and bioinformatics analysis showed the putative binding site of miR-126 and S1PR2. Considering the characteristics miRNAs function, we hypothesized that the role of miR-126 in IBD was mediated by the interactive endogenous relation between miR-126 and S1PR2.

Firstly, the results of luciferase reporter gene assay supported that S1PR2 was a direct target of miR-126 in Caco-2 cells. The overexpression or knockdown of miR-126 directly affected the expression of S1PR2 protein *in vitro.* Intestinal mucosal barrier dysfunction caused by changes in cell permeability and intracellular junction is a key factor in the pathogenesis of IBD. Tight junction between intestinal epithelial cells is the complex consisting of a series of proteins and fats, which exists widely between cells and cells. The tight junction proteins, the main component of tight junctions, are important for tight junctions to maintain the appropriate structure and function, including Occludin, E-Cadherin, ZO-1. In enterocyte barrier injury model of Caco-2 cells induced by LPS, S1P contributed to stabilize the intestinal barrier function via modulation of SP1R2. However, in the present study, it was strongly confirmed that miR-126 overexpression contributed to intestinal mucosal barrier dysfunction as shown by abolishment of the protection effect of S1P for the integrity and permeability of intestinal mucosal barrier.

S1P-activated SP1R2 could promote cell migration via modulating PI3K/AKT signaling pathway. A great deal of researches showed that PI3K/AKT signaling pathway was an important regulator for intestinal mucosal barrier function in intestinal mucosa injuries. We revealed that S1P/S1PR2 protected intestinal mucosal barrier function via a PI3K/Akt dependent mechanism *in vitro.* Based on the above results, we further validated the role of miR-126 in the regulation mechanism of S1P/S1PR2/PI3K/AKT signaling pathway for intestinal barrier function. MiR-126 silencing possessed obvious protective effects on the intestinal barrier function.

In conclusion, our results confirmed that miR-126 may serve as a promotion factor for the progress of IBD via targeted modulation of S1PR2. The study provided direct evidence that miR-126 down-regulated S1PR2 and then prevented the activation of PI3K/AKT signaling pathway, which ultimately could damage intestinal mucosal barrier function. Accordingly, miR-126 will be a new therapeutic target for IBD.

## Acknowledgments

This work was part of the Program on Wenzhou Science & Technology Bureau, No: Y20160026.

This study was funded under Zhejiang Provincial Natural Science Foundation, grant number LY17H030010.

